# Soil texture shapes biochar-induced shifts in microbial communities and severity of potato common scab

**DOI:** 10.1101/2025.07.11.664502

**Authors:** Alena Maslova, Jan Kopecky, Vaclav Tejnecky, Filip Mercl, Marketa Sagova-Mareckova

## Abstract

Biochar is widely recognized for its potential to enhance soil carbon, yet its influence on soil chemistry, microbial communities, and plant disease dynamics remains uncertain. This study explored how biochar affects the severity of common scab, a globally significant potato disease, through its interactions with soil properties and microbial communities. Two contrasting soils planted with potatoes were amended with fresh biochar and biochar aged within the soils for one and two years. Bulk and tuberosphere soil samples were analyzed for nutrient and micronutrient availability, microbial community composition using 16S rRNA gene amplicon sequencing and basal respiration. In sandy soil V, biochar induced immediate but transient shifts in soil chemistry and microbial structure, while in silty soil Z, changes emerged gradually and intensified with biochar aging. These dynamics mirrored the progression of common scab severity, suggesting a strong link between nutrient availability, microbial community shifts, and disease outcomes. The contrasting responses of the two soils were attributed to differences in cation exchange capacity and particle size, which influenced nutrient retention and microbial community structure, including populations of pathogenic *Streptomyces*. Notably, the exchangeable surface area of soil particles emerged as a factor modulating biochar effects on disease severity. These findings highlight the importance of considering soil texture and microbial composition when applying biochar, emphasizing that site-specific characteristics can critically shape biochar’s impact on plant health.

**Importance:** Biochar is increasingly used to improve soil quality and health, but its effects on plant diseases are not fully understood. This study showed that biochar can either reduce or worsen common scab of potatoes depending on soil type. By examining how biochar changes soil chemistry, nutrient availability and microbial communities, we found that soil texture and nutrient exchange capacity influence the disease outcomes. In silty soil, biochar decreased nutrient availability and promoted pathogenic bacteria that cause common scab, while in sandy soil, it did not. These findings highlight the need to consider local soil conditions before applying biochar in agriculture. Understanding how soil properties influence microbial responses helps farmers and researchers make better decisions to protect crops and improve soil health.

## Introduction

Potato common scab (CS), caused by phytopathogenic Streptomyces spp., is a globally widespread disease leading to losses in crop yields with significant economic impacts (1). The disease management remains challenging due to the intricate interplay between soil properties, microbial communities, plant genotype, and pathogen virulence (2–4). In addition, disease severity is modulated by multiple soil factors, including pH, moisture, nutrient availability, and microbial community composition (5, 6), underscoring the need for integrated approaches that consider both biotic and abiotic soil components (2, 5, 6).

Biochar, a carbon-rich byproduct of biomass pyrolysis, has emerged as a promising soil amendment due to its ability to enhance soil structure, promote carbon sequestration, increase nutrient retention, and modulate microbial communities (7, 8). Its high surface area, alkaline nature, and capacity for cation exchange and adsorption of organic molecules make it a potent modifier of soil chemical and biological processes (9, 10). As biochar ages, it is incorporated into soil aggregates, while promoting the stabilization of rhizodeposits and microbial products, which further influences its performance in soil (11).

Biochar has been shown to influence plant–microbe–pathogen interactions by altering nutrient dynamics, microbial signaling, and the abundance of plant growth-promoting rhizobacteria (PGPR) (12, 13). However, its effects on soil-borne pathogens remain inconsistent and context-dependent, often varying with soil texture, mineralogy, and microbial baseline conditions (14, 15). In addition, biochar influences the availability of micronutrients, cation exchange capacity and adsorption processes, which may also influence the outcomes of biochar application (12, 16).

Possible mechanisms by which biochar can induce plant disease resistance include increased nutrient content or supply, pH, reduced bioavailability of phytotoxins, and improved soil physical characteristics (13). The modified microbial community may change soil-born pathogen pressure either by competition for nutrients and space, or through production of pathogen-inhibiting compounds. More complex processes include also adsorption of microbial signaling molecules, enzymes and toxic metabolites, thus reducing the concentration of virulence factors in the root zone (9, 17).

Finally, biochar amendments to soil can stimulate plant growth promoting bacteria, which suppress the disease either directly by antibiotics against the pathogen or indirectly by increased plant fitness (12). Yet, despite the growing interest in biochar applications, few studies have simultaneously examined its long-term effects on soil chemical properties, microbial community structure, and disease outcomes in field-relevant systems. Moreover, the role of soil texture and chemistry in modulating these effects remains poorly understood, particularly in relation to diseases and soil-born pathogens (14).

In this study, we investigated how biochar application affects common scab severity, soil chemistry, and microbial communities in two contrasting soils differing in texture and CEC. We followed changes over a three-year period of biochar aging to capture both immediate and aging-related effects. We focused on the following hypotheses: 1) Soil texture and CEC modulate biochar effects, with finer-textured, high-CEC soils exhibiting stronger chemical and microbial responses. 2) Long-term effects of biochar differ from short-term responses, due to aging-related changes in biochar reactivity and microbial adaptation. 3) Biochar suppresses CS severity by enhancing nutrient availability and supporting beneficial microbial taxa, particularly PGPR.

## Results

### Soil chemical characteristics

In the beginning of the experiment, soils from both sites (V and Z) had the same carbon content of 1.6% dry weight (dw) and also the differences in extractable Fe, P, N and S content were not significant. K_a_ and K_e_ contents (K extractable by ammonium acetate or EDTA) were significantly higher in soil V and Ca_a_ and Ca_e_ contents in soil Z. The soils differed in the proportion of clay, silt, and sand (18). However, the differences between the two soils changed after biochar application. They differed significantly in P, Fe_e_, Fe_a_, Ca and S (but not K) after one year and in Fe_e_, Ca_a_ and S after two years from application (Table S2).

Biochar application caused an increase of carbon content, which almost did not change over the years of aging (Tables 2, S3, Fig. S1). Nitrogen content did not change after biochar application in either soil, only in Z soil a significant decrease occurred in two years old compared to the freshly added biochar. P_e_ content in V soil significantly decreased after one year, while in Z soil significantly increased after one and two years. Both K_a_ and K_e_ in V soil significantly decreased between fresh application and after two years, while in Z soil both K fractions significantly increased over time. Ca_a_ increased in V soil after one year, and Ca_e_ after two years, while in Z soil both forms of Ca were significantly higher in both years after biochar application. Fe_a_ in soil V increased after two years while Fe_e_ did not change after biochar application, but in Z soil the increase was significant after both years for Fe_a_ and after the first year for Fe_e_. The S content did not significantly change in either soil after biochar application. The original soil pH was slightly acidic in both soils (V, Z) and the biochar application caused a significant increase in both soils one and two years after application (Tables 2, S3, Fig. S1).

### Potato leaf element content

N and K contents in leaves did not differ between the two soils and also did not respond to any of the treatments although there was a decreasing trend in N after biochar additions. Fe content increased significantly in Z soil one year after application (Fig. S2, Table S4).

### Common scab severity

After fresh biochar application CS severity remained unchanged in both soils but it significantly increased in soil Z one and two years after biochar application (Fig. 1, Table S3). The observation of increased CS severity was further confirmed by increased proportion of potentially pathogenic streptomycetes in the community (Fig. S3). In bulk soil at the start of the experiment, V soil had a higher proportion of potentially pathogenic streptomycetes than Z soil, and the proportion remained similar after biochar addition. In tuberosphere, the proportion was higher in both soils and the group of *S. niveiscabiei* appeared in the community. The two soils differed in reaction of potentially pathogenic streptomycetes to biochar 1 and 2 years after application, while in V soil the proportion remained the same, it increased in Z soil in both years.

**Fig. 1.**
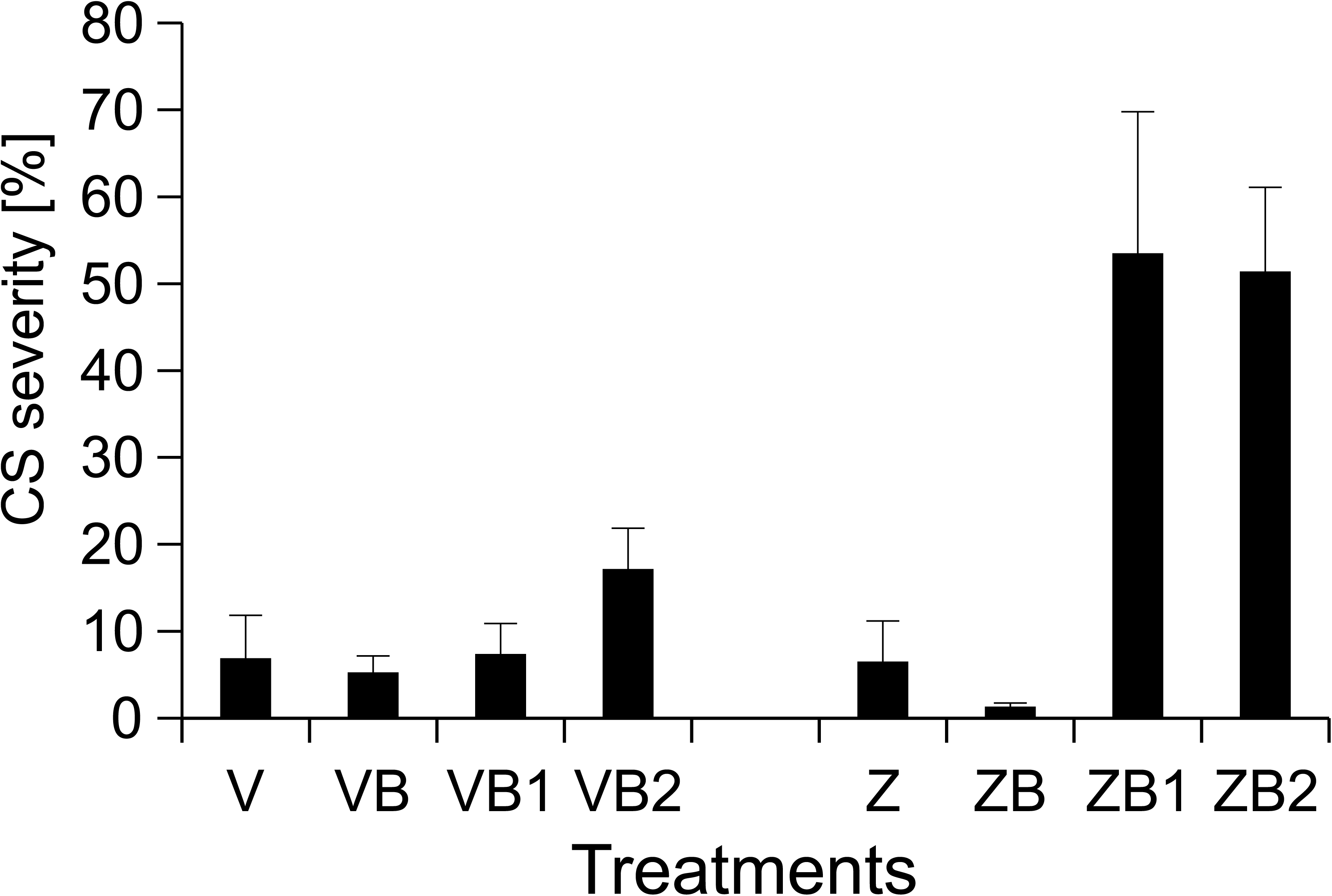
The severity of common scab in potatoes grown in untreated soils V and Z, soils with freshly added biochar (B), 1 year (B1), and 2 years (B2) after biochar application. Disease evaluation in percent of the covered tuber surface area (43). Letters indicate statistical significance (Dunn’s test, p < 0.05) between treatments (lower case - site V and upper case - site Z).

### Soil respiration

The respiration rate was similar in both soils Z (143.8 ± 18.6 nmol O_2_ h^-1^ g^-1^) and V (96.5 ± 16.2 nmol O_2_ h^-1^ g^-1^). However, in soil V the respiration rate significantly decreased in the second year after biochar application, while in soil Z, the decrease occurred after one year and remained low in the second year (Fig. 2, Table S3).

**Fig. 2.**
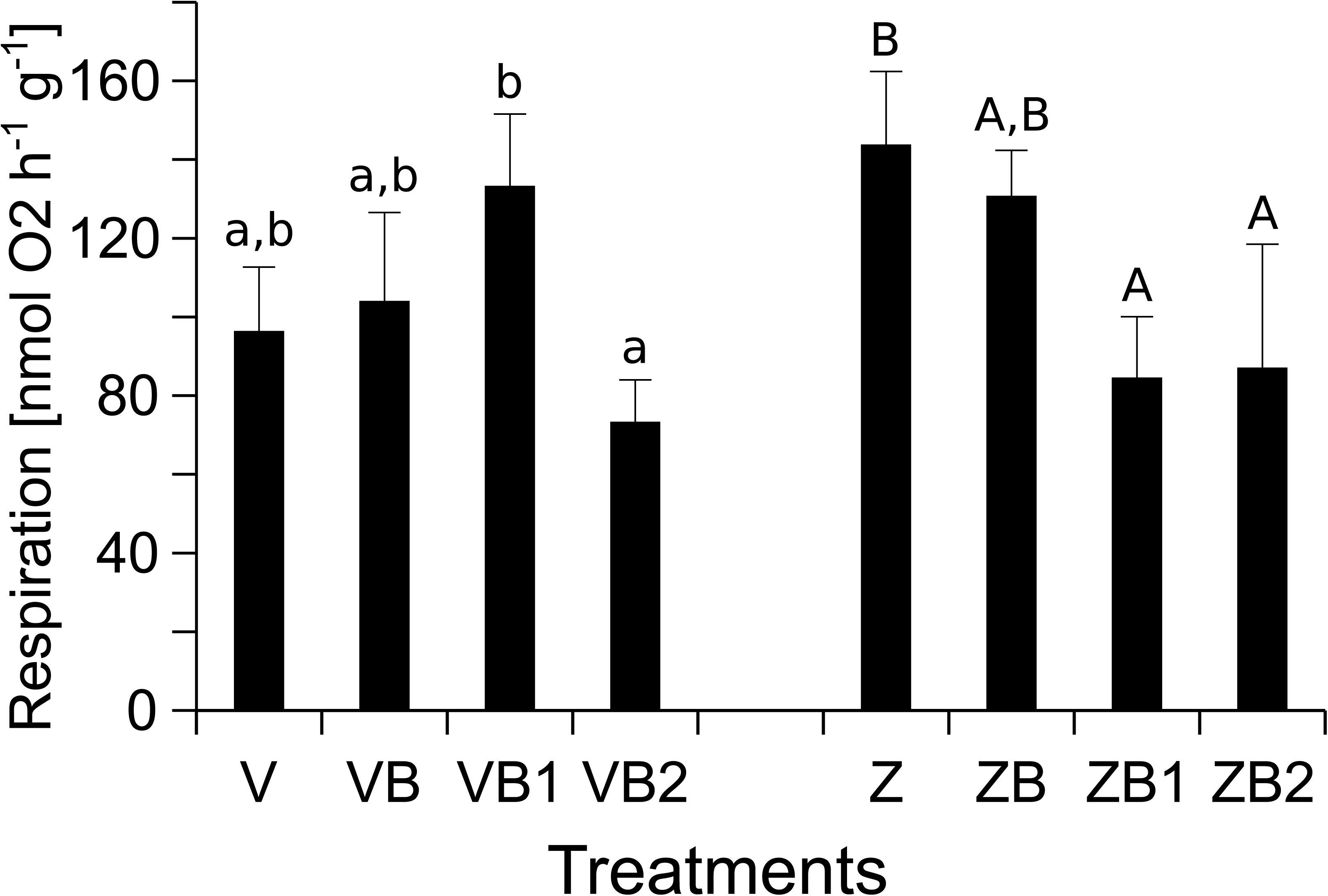
Respiration rate in untreated soils V and Z, soils with freshly added biochar (B), 1 year (B1), and 2 years (B2) after biochar application. Letters indicate statistical significance (Dunn’s test, p < 0.05) between treatments (lower case - site V and upper case - site Z).

### Prokaryotic communities

The non-metric multidimensional scaling (NMDS) plot demonstrated shifts of prokaryotic communities after biochar additions. Prokaryotic communities significantly differed between the two original soils and also reacted differently to biochar additions in tuberosphere (Fig. S4, Table S5).

At the start in soil V, prokaryotic communities in the untreated control and soil with freshly added biochar (fresh soils) differed from those after 1 and 2 years but significant difference occurred also between communities from soils after 1 and 2 years from application. In soil Z, the separation was similar, only prokaryotic communities of fresh soils were more dispersed, and a significant shift occurred between the fresh soils and soil after 1 year (Table S5). In tuberosphere soil, the separation of prokaryotic communities in the two soils was similar. In tuberosphere soil V, a significant difference in prokaryotic community occurred between fresh soils and addition after 1 year and between 1 and 2 years after addition. In soil Z, significant differences occurred between the fresh soils and the first year after addition (Fig. 3, Table S5). Prokaryotic communities were differently related to soil chemical properties in the two tuberosphere soils. In soil V, prokaryotic communities were significantly related to Ca content and soil respiration rate one year after application, while they were related to soil pH after 1 and 2 years of application, and to Fe_a_ content after two years of application. The highest diversity correlated with control soils (Fig. 3a). In soil Z, prokaryotic communities were more clearly separated by year of biochar application and the relationships to soil chemistry were stronger than in soil V. In particular, in the control soil without biochar prokaryotic communities were related to soil respiration rate, 1 year after biochar application to soil pH, Ca_e_, P_e_, K_e_, Fe_e_ contents and Simpson diversity indices, and 2 years after biochar application to Fe_a_, K_a_, and Ca_a_ contents (Fig. 3b). The prokaryotic community composition did not correlate with CS severity in either of the soils (Fig. 3a,b).

**Fig. 3.**
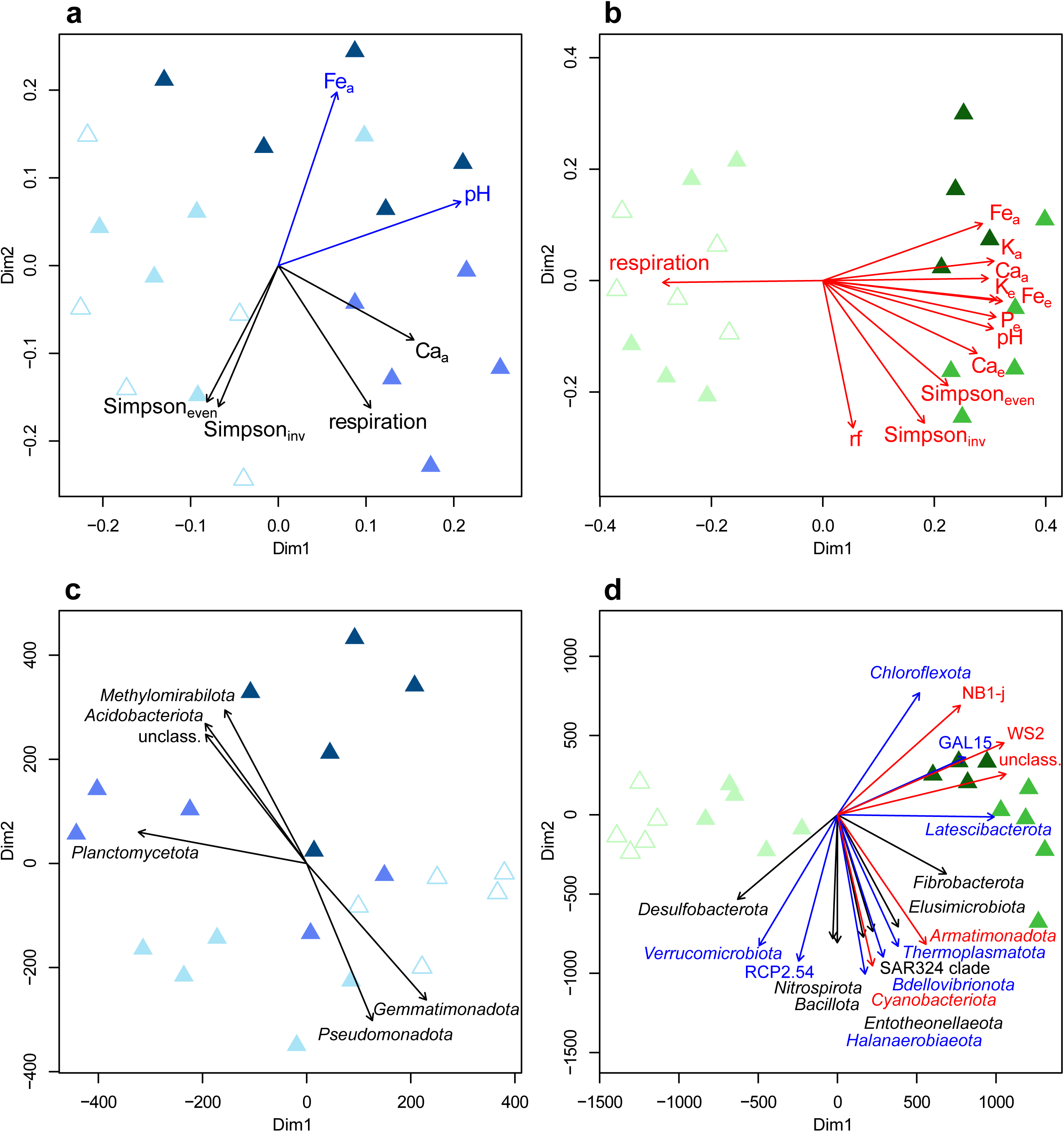
Comparison of prokaryotic community composition (A, B) and soil chemical properties (C, D) in the tuberosphere of potato grown in soil V (A, C) and Z (B, D), and their correlation with soil chemical properties, respiration rates and diversity measures (A, B), and with relative abundances of selected prokaryotic taxa (C, D), respectively. Symbols (triangles) represent prokaryotic communities in untreated soil (open symbols), soil with freshly added biochar (light color), 1 year (medium-dark), and 2 years after biochar application (dark), and vectors show the correlated parameters. Non-metric multidimensional scaling based on Bray-Curtis distance matrix. Color of vectors indicates statistical significance of observed relationships P ≤ 0.05 (black), P ≤ 0.01 (blue) and P ≤ 0.001 (red). Abbreviations: rf - area below rarefaction curve; a, e in subscript - extraction with ammonium acetate and EDTA, respectively; Simpson_inv_ – inverse Simpson index; Simpson_even_ – Simpson evenness index, unclass. – unclassified bacteria.

In the NMDS plots of soil chemical properties, in soil V (Fig. 3c) the fresh soils, control and soil after addition of fresh biochar, differed only weakly from the soil one and two years after biochar application, while in soil Z (Fig. 3d) all treatments were clearly separated from each other. Chemical properties of fresh soils V were significantly correlated to the relative abundance of *Pseudomonadota* and *Gemmatimonadota*, soil V after one year to *Planctomycetota* and after two years to *Methylomirabilota* and *Acidobacteriota* (Fig. 3c). In soil Z, soil chemical properties one year after biochar application were related to the increased proportion of *Latescibacterota* and *Fibrobacterota*, and after the second year they were related to *Chloroflexota*, and clades NB1-j, WS2 and GAL15 (Fig. 3d). Many phyla showed negative correlation to chemical changes occurring in the second year after biochar addition, e.g. *Cyanobacteriota, Armatimonadota, Verrucomicrobiota, Halanaerobiaeota, Bdellovibrionota, Thermoplasmatota,* and clade RCP2.54 (Fig. 3d).

In both soils at the start, a higher proportion of *Actinomycetota* was observed compared to the end in the tuberosphere, where a higher proportion of *Acidobacteriota*, *Verrucomicrobiota* and *Methylomirabilota* occurred (Fig S5A). *Bacteroidota* appeared in tuberosphere of both soils while at the start they were more represented in soil Z than soil V. In soil V, proportion of *Actinomycetota* and *Methylomirabilota* increased after the 1 and 2 years after biochar application, and at the same time a decrease in the proportion of *Pseudomonadota* was observed. In tuberosphere of soil V, *Acidobacteriota* and *Methylomirabilota* increased one year after the biochar application, *Pseudomonadota* gradually decreased, and *Actinomycetota* decreased after biochar application, but restored the original proportion two years later. In soil Z, *Actinomycetota* increased after the biochar application and dropped in the following years, and the proportion of *Acidobacteriota, Chloroflexota* and *Planctomycetota* also increased. The proportion of *Actinomycetota*, *Chloroflexota* and *Planctomycetota* in tuberosphere soil Z increased after biochar treatment, while *Verrucomicrobiota, Acidobacteriota, Bacteroidota*, and *Thermoproteota* decreased (Fig. S5A).

Within the phylum *Actinomycetota* in soil V, biochar application increased the proportion of orders *Gaiellales*, *Soilrubrobacterales*, *Microtrichales*, and clades MB-A2-108 inc. sedis and IMCC26256, and decreased the proportion of orders *Micrococcales* and *Propionibacteriales*. An increased proportion of *Micrococcales* occurred 2 years after the biochar application in the tuberosphere soil V, where the proportion of MB-A2-108 also increased, while *Frankiales* decreased. In soil Z, the proportion of orders *Gaiellales*, *Soilrubrobacterales*, *Propionibacteriales* and *Frankiales* decreased and the proportion of *Microtrichales* and MB-A2-108 increased after the biochar application. In the tuberosphere soil Z, the proportion of *Gaiellales* and *Corynebacteriales* decreased and the proportion of *Micrococcales*, *Soilrubrobacterales*, *Propionibacteriales*, *Microtrichales*, MB-A2-108, *Frankiales*, *Streptomycetales* and *Pseudonocardiales* increased after the biochar application (Fig. S5B). Within the phylum *Pseudomonadota*, the proportion of order *Rhizobiales* was higher in the tuberosphere of both soils than in the bulk soil. In both the tuberosphere and bulk soil V, the proportion of orders *Burkholderiales*, *Sphingomonadales* and *Xanthomonadales* decreased after biochar application. In the bulk soil Z, the proportion of orders *Rhizobiales* and *Sphingomonadales* first decreased and then increased above the initial level in the second year after the application. One and two years after the biochar application in bulk soil Z, the proportion of *Rhodobacterales* and *Sphingomonadales* increased, while the proportion of *Rhizobiales* in the tuberosphere decreased after biochar application (Fig. S5C).

Changes in prokaryotic community composition compared at the level of individual ASVs followed the patterns described at the order level (Fig. 4,). Biochar addition increased numbers of ASVs in both soils but the effect diminished with aging. The most distinct difference between treatments with old biochar and fresh soils was the proportion of ASVs belonging to *Pseudomonadota* dominating at the start as well as at the end of the experiment in tuberosphere with freshly added biochar compared to domination of those from *Actinomycetota and Chloroflexota* in soils aged 1 and 2 years after biochar additions (Figs. S6-9).

**Fig. 4.**
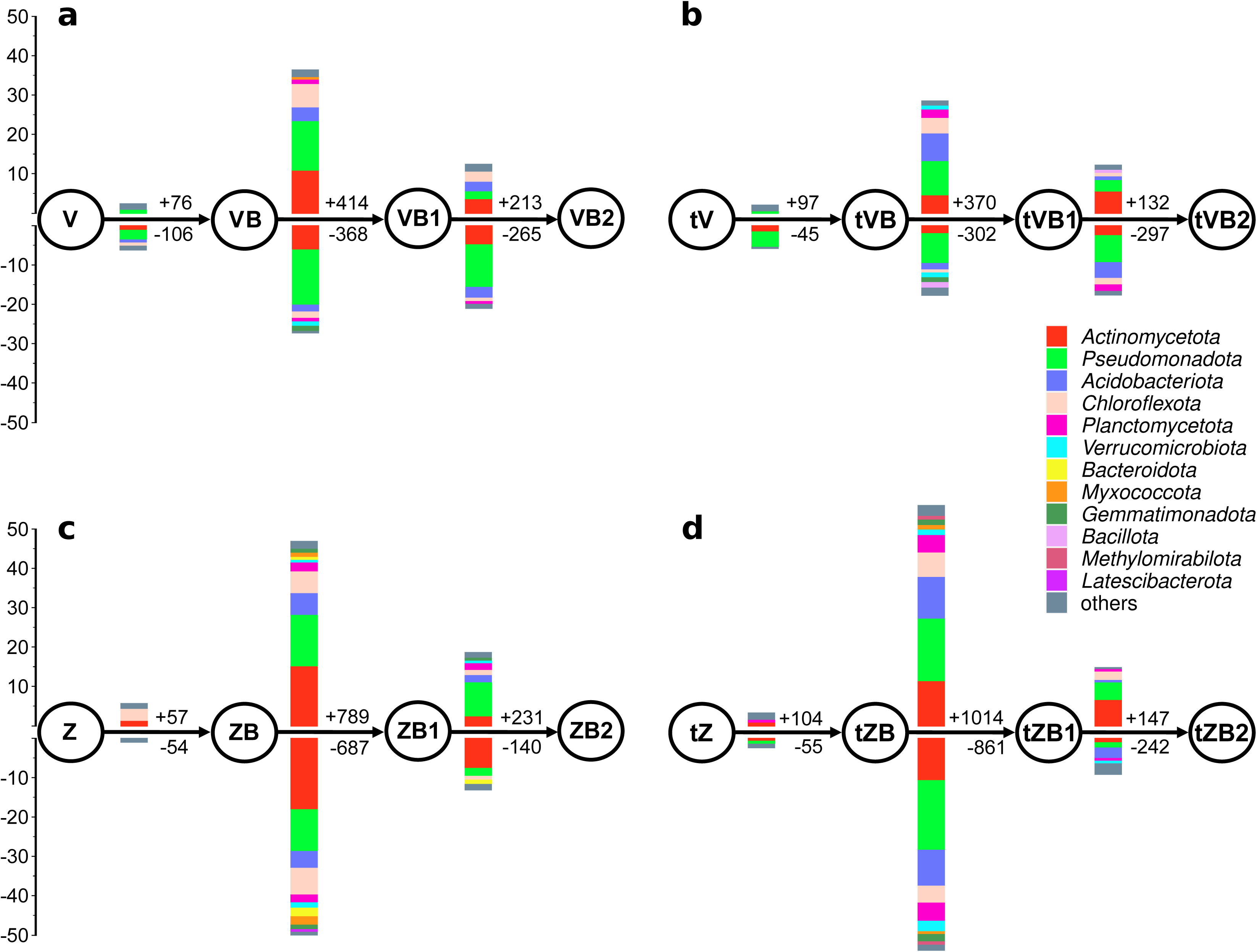
The numbers of ASVs in bulk and potato tuberosphere soil at the sites V (A, B) and Z (C, D),, that contribute significantly to the differences between prokaryotic communities in untreated soils (V,Z or tV, tZ), soils with freshly added biochar (B), 1 year (B1), and 2 years (B2) after biochar application. The plus and minus signs indicate numbers of ASVs increasing and decreasing their proportion in direction of the arrow. The barplots show the proportions of those ASVs grouped at phylum level (Metastats, *p* < 0.05).

### Bacterial diversity

Prokaryotic diversity was affected by biochar additions. At the start in bulk soil V, both inverse Simpson and Simpson evenness indices were significantly increased 1 and 2 years after application, while in bulk soil Z, a significant increase occurred only in the second year. In tuberosphere in soil V, there was a significant decrease of inverse Simpson index in the second year, while in soil Z a significant increase of both indices occurred after one year from biochar application (Table S5).

### Predicted metabolic pathways

Effects of biochar treatment on functions of prokaryotic communities were estimated based on the 16S rRNA gene community profiles. The major difference between sites was found in central fermentation pathways with higher proportion in bulk soil of Z and tuberosphere soil of V compared to the other treatments (not shown). The most significant result was for hydrogenotrophic methanogenesis, in which again Z soil had higher proportion of representative ASVs in bulk but also in tuberosphere soil except after 2 years from biochar application compared to the other treatments (Fig. S10).

## Discussion

Biochar application did not significantly change CS severity in soil V but increased it in soil Z. That was in contrast to our expectation and previous findings in various patho-systems as plant diseases were mostly reduced after biochar application (14, 15, 19).

The suppression of CS by biochar has previously been attributed to enhanced soil porosity and water retention (14). In other patho-systems, mechanisms such as increased dissolved carbon, improved nutrient availability, shifts in microbial community structure, and induced systemic resistance have been proposed (15, 19). In our study, biochar application elevated soil carbon and pH in both soils, yet triggered distinct chemical changes. In soil V, potassium levels declined while calcium and iron increased. Conversely, in soil Z, phosphorus, potassium, calcium, and iron all rose, while nitrogen decreased. These contrasting responses are largely explained by soil texture: soil V is sandy and lacks fine particles, whereas soil Z contains 60% silt. While clay content is often considered the dominant factor in soil chemical reactivity (20, 21), our findings suggest that silt may play a more critical role in element release (22). Multiple physico-chemical mechanisms are likely at play. Notably, in the finer soil Z, small biochar particles increased bulk density and filled pore spaces (14, 23, 24), potentially altering water retention. This aligns with previous observations that both overly wet and overly dry soils can increase CS severity (25, 26). Our analysis of predicted metabolic pathways further supports this, revealing a correlation between increased abundance of obligate anaerobic methanogens and higher disease severity—indicative of reduced oxygen availability, as also noted by Longepierre et al. (24).

Secondly, the high proportion of fine silt particles in soil Z suggests a significantly larger exchange surface compared to the sandy soil V, which likely competes with biochar surfaces for cation adsorption. Biochar is known for its high surface charge density, enabling effective cation retention through cation exchange. Its large surface area, internal porosity, and diverse surface chemistry—comprising both polar and non-polar sites—also facilitate the adsorption of organic molecules and associated nutrients (19, 27, 28). In most studies, enhanced cation exchange capacity (CEC) has been identified as a key mechanism behind reduced nutrient leaching following biochar application (23, 29). However, some findings suggest that this effect may diminish over time; for instance, Jones et al. (27) reported a decline in biochar-induced CEC after three years, with no sustained improvement in nutrient availability. In our study, the silt-rich soil Z, with a CEC of approximately 20, likely retained a higher proportion of cations introduced via biochar compared to the sandy soil V, which had a CEC of around 6 (30). This greater retention capacity may contribute to nutrient availability and aligns with earlier observations linking increased CEC to higher CS severity (31).

Finally, the contrasting nutrient dynamics between the two soils were also reflected in plant tissue composition. In plants grown in soil Z, iron content increased significantly, while no notable changes were observed in plants from soil V. This differential response further underscores the role of exchangeable surfaces and CEC in mediating iron availability, potentially through immobilization mechanisms that enhance plant uptake (21, 32). Moreover, the rise in pH following biochar application altered the composition and relative abundance of pathogenic *Streptomyces* spp., particularly in soil Z. These shifts may have influenced CS severity by modulating the virulence of specific strains, as previously suggested by Loria et al. (33).

Prokaryotic community composition corresponded to chemical trends in both soils because in soil V, the differences were rather small and gradual, while in soil Z, clearly distinguished communities occurred between the soil freshly added biochar on one side and soils aged with biochar after one and two years on the other side. More specifically, chemical conditions in soil V were reflected in fresh soils by increased *Pseudomonadota* and *Gemmatimonadota*, and in aged biochar soils by increased *Vicinamibacterales* (*Acidobacteriota*), *Planctomycetota* and *Methylomirabilota*. All of those taxa were previously reported as responding to biochar additions but partially in an opposite way. *Acidobacteriota* were rather seen in oligotrophic conditions without biochar (34), while *Pseudomonadota* increased with added carbon substrates. However, in another study (35) *Pseudomonadota* also decreased after biochar additions. In soil Z, only *Rhizobiales*, possibly *Chthoniobacterales* were enriched in fresh soil, but biochar addition induced an increase in many taxa including *Latescibacterota*, *Fibrobacterota*, WS2, GAL15 and *Burkholderiales* in soils with aged biochar. Thus, in both soils, the changes included typical plant growth promoting taxa, mostly decreasing (*Rhizobiales, Sphinomonadales, Xanthomonadales*) but also increasing (*Burkholderiales*) with biochar additions. Changes in PGPR with biochar were determined previously (12) but it was observed that specific soil properties may diminish the positive effect of PGPR bacteria because biochar may adsorb signaling molecules or antibiotics, which play role in disease suppressivity and prokaryotic community interactions (9).

The relative abundance of *Pseudomonadota* decreased in soil V after the biochar addition, which corresponded to the decrease of *Pseudomonadota* in a previous study (36). In comparison, the proportion of *Actinomycetota* increased after the addition of biochar in bulk soil V and in the tuberosphere of both soils, while there was a decrease in the bulk soil Z. Similarly, the biochar application increased *Actinomycetota* proportions previously (37, 38). On one hand, the increase of *Actinomycetota* may indicate an enhancement of *Streptomyces* pathogens, which belong to *Actinomycetota*, on the other hand it may be connected to pathogen antagonistic activities provided by non-pathogenic streptomycetes (39, 40). It seemed that in our study, particularly the relative proportion of pathogenic streptomycetes increased in potato tuberosphere, and in soil Z, where CS severity increased significantly, also in response to biochar addition. That is further supported by significant change in the pathogenic community composition and proportion suggesting different aggressivity of pathogens (33) as well as successful competition of pathogens for DOC (41). Similarly to changes in prokaryotic community composition, there were changes in prokaryotic diversity leading to diversity decrease in soil V and increase in soil Z. In our study these changes seemed that similarly to the previous findings (42) prokaryotic community responded to new sources of DOC, which occurred mostly in soil Z.

## Conclusion

Our findings revealed that the impact of biochar on soil microbial communities and common scab severity is highly soil-dependent. Biochar-induced shifts in soil chemistry were closely aligned with changes in prokaryotic community structure and CS disease outcomes.

Notably, soil particle size and associated exchangeable surface area and cation exchange capacity (CEC) emerged as factors influencing (micro)nutrient dynamics and microbial responses. These physicochemical properties affected nutrient availability, plant nutrient uptake, and the composition of pathogenic Streptomyces populations. In silty soil, biochar favored the emergence of more aggressive streptomycete taxa, while in sandy soil, such shifts were absent. These results underscore the importance of considering soil texture, CEC, and the local pathogen community when applying biochar in agricultural systems. Optimizing biochar use based on these parameters may help mitigate unintended increases in soil-borne disease severity and promote more predictable outcomes for soil health and plant productivity.

## Materials and methods

### Sites

The soils came from two sites Vyklantice (V) and Zdirec (Z) located in the Czech Moravian Highlands, which we have studied previously. The soils V and Z differed in soil texture (clay : silt : sand : gravel [%] 0:30:62:8 vs. 6:56:35:3), cation exchange capacity (CEC) (6.0 vs. 20.1 cmol kg^-1^) and pH (6.30 vs. 6.98), but did not significantly differ in pathogen quantities determined by qPCR of thaxtomin biosynthetic gene (soil V: 9.8 × 10^3^, soil Z: 8.8 × 10^4^ *txtB* gene copies g^-1^ (4, 18).

### Biochar

Biogenic waste (grain husks, sunflower peels, and fruit pulp) was used as a raw material to produce biochar. The raw material was heated to a temperature of 500-600 °C during pyrolysis (Sonnenerde, Riedlingsdorf, Germany). The resulting biochar was of fine dust-like particles and was enriched particularly in Ca, K, P and S (Table S1). The biochar was handled in clean environment always with gloves, so no significant additions of microbes with biochar were expected.

### Experimental design

The experiment started by adding biochar to 12-L perforated pots with soil (5 replicate pots of each soil) in the spring of 2014, 2015, and 2016. Biochar in the amount of 0.13 kg per pot (corresponding to 2 kg of biochar homogeneously distributed in 20 cm topsoil per m^2^) was carefully mixed with homogenized fresh soil collected on site and kept buried to soil under tree shade till the beginning of potato cultivation experiment. The goal was to compare the effect of biochar additions over three years in soil with and without potato plants.

The experiment was set up in May 2016, five replicate pots of each soil and treatment, i.e. the pots with added biochar in each year and the pots with untreated control soils (Table 1), were planted with Agria, a CS susceptible potato cultivar, and kept sunk in the field, soil up to the rim without irrigation or fertilization but receiving regular pesticide treatment throughout the growing season. Samples of bulk soil were collected before potato planting. The samples of tuberosphere soil, plant leaves and potatoes were collected 90 days after planting, when the disease is fully developed but the plant is still living, thus actively modifying its microbiome. Tuberosphere soil samples of approximately 30 mL were taken carefully no further than 3 mm from the tuber using a sterile spoon, filled to 50 mL falcon tubes, transferred to laboratory at −18 °C, and stored at −80 °C before the analysis (for details see 18).

**Table 1.**
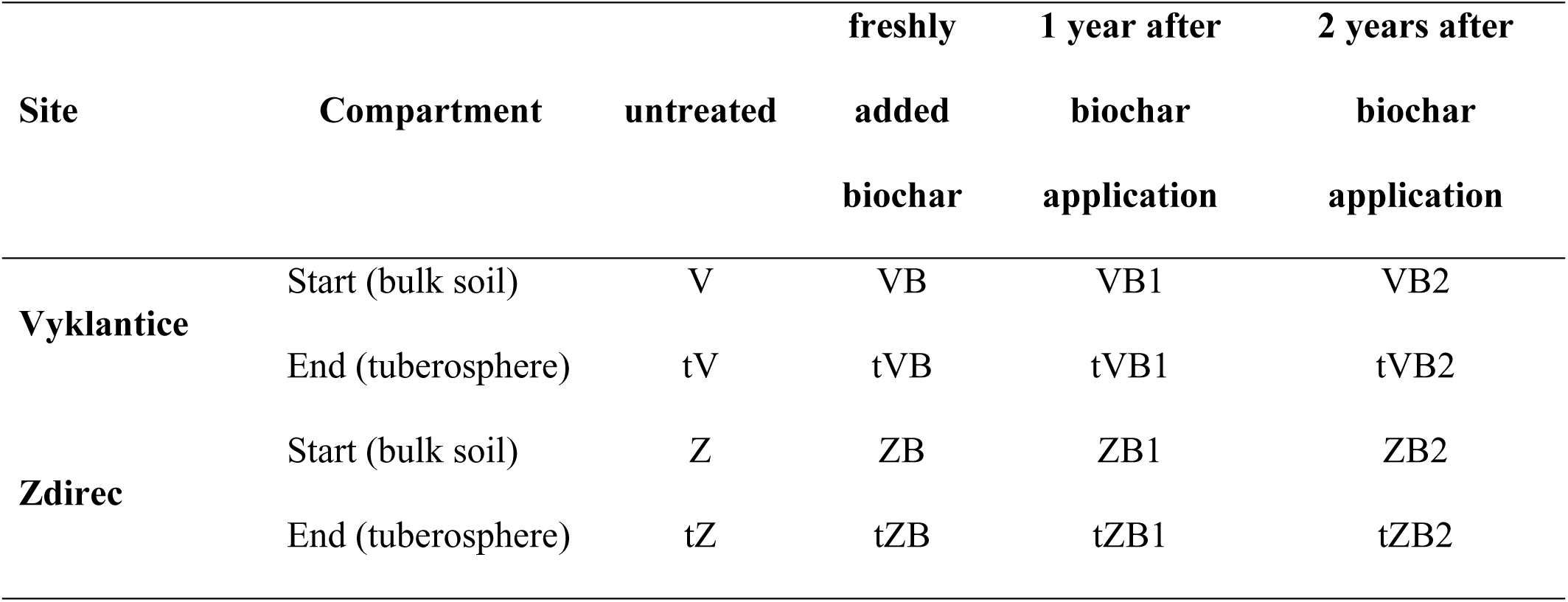
Soil treatments and designation.

**Table 2.** Changes in C, Fe, K, Ca, P, N and S contents and soil pH after biochar application (0, 1, and 2 years). Abbreviations: a, e in subscript indicate extraction with ammonium acetate and EDTA. Superscript letters indicate statistical significance (Dunn’s test, p < 0.05) between treatments (lower case - site V and upper case - site Z).

|  | V | VB | VB1 | VB2 |
| --- | --- | --- | --- | --- |
| **pH** | $6.37 \pm 0.12^a$ | $6.52 \pm 0.17^{a,b}$ | $6.89 \pm 0.11^c$ | $6.70 \pm 0.19^{b,c}$ |
| **C [%]** | $1.61 \pm 0.24$ | $2.09 \pm 0.22$ | $2.14 \pm 0.19$ | $2.03 \pm 0.32$ |
| **N [%]** | $0.144 \pm 0.040$ | $0.180 \pm 0.020$ | $0.180 \pm 0.020$ | $0.166 \pm 0.010$ |
| **S [%]** | $0.040 \pm 0.004$ | $0.043 \pm 0.004$ | $0.045 \pm 0.017$ | $0.044 \pm 0.008$ |
| **Fe<sub>e</sub> [mg kg<sup>-1</sup>]** | $301 \pm 10$ | $300 \pm 29$ | $295 \pm 21$ | $304 \pm 30$ |
| **K<sub>e</sub> [mg kg<sup>-1</sup>]** | $222 \pm 26^{a,b}$ | $289 \pm 15^b$ | $223 \pm 12^{a,b}$ | $176 \pm 13^a$ |
| **Ca<sub>e</sub> [mg kg<sup>-1</sup>]** | $1450 \pm 63^a$ | $1572 \pm 81^{a,b}$ | $1606 \pm 102^{a,b}$ | $1680 \pm 119^b$ |
| **P<sub>e</sub> [mg kg<sup>-1</sup>]** | $71.8 \pm 2.6^{a,b}$ | $85.0 \pm 7.8^b$ | $68.2 \pm 4.9^a$ | $74.3 \pm 5.3^{a,b}$ |
| **Fe<sub>a</sub> [mg kg<sup>-1</sup>]** | $0.40 \pm 0.14^a$ | $0.30 \pm 0^a$ | $0.34 \pm 0.09^a$ | $1.64 \pm 0.21^b$ |
| **K<sub>a</sub> [mg kg<sup>-1</sup>]** | $234 \pm 25^{a,b}$ | $317 \pm 27^c$ | $265 \pm 26^{b,c}$ | $194 \pm 24^a$ |
| **Ca<sub>a</sub> [mg kg<sup>-1</sup>]** | $1398 \pm 28^a$ | $1510 \pm 64^{a,b}$ | $1614 \pm 119^b$ | $1440 \pm 103^{a,b}$ |

B. Site Z
|  | Z | ZB | ZB1 | ZB2 |
| --- | --- | --- | --- | --- |
| **pH** | $6.14 \pm 0.13^A$ | $6.34 \pm 0.05^{A,B}$ | $7.33 \pm 0.05^C$ | $7.15 \pm 0.19^{B,C}$ |
| **C [%]** | $1.60 \pm 0.14^A$ | $2.48 \pm 0.23^B$ | $2.04 \pm 0.11^{A,B}$ | $1.98 \pm 0.27^{A,B}$ |
| **N [%]** | $0.176 \pm 0.020^{A,B}$ | $0.194 \pm 0.010^B$ | $0.168 \pm 0.010^{A,B}$ | $0.148 \pm 0.030^A$ |
| **S [%]** | $0.027 \pm 0.005$ | $0.026 \pm 0.002$ | $0.025 \pm 0.002$ | $0.025 \pm 0.004$ |
| **Fe<sub>e</sub> [mg kg<sup>-1</sup>]** | $389 \pm 30^A$ | $395 \pm 48^A$ | $745 \pm 44^B$ | $706 \pm 25^{A,B}$ |
| **K<sub>e</sub>** [mg kg<sup>-1</sup>] | 68.3 ± 11<sup>A</sup> | 105 ± 7<sup>A,B</sup> | 196 ± 9<sup>C</sup> | 184 ± 25<sup>B,C</sup> |
| **Ca<sub>e</sub>** [mg kg<sup>-1</sup>] | 1798 ± 107<sup>A</sup> | 2060 ± 158<sup>A,B</sup> | 2896 ± 175<sup>C</sup> | 2488 ± 71<sup>B,C</sup> |
| **P<sub>e</sub>** [mg kg<sup>-1</sup>] | 17.1 ± 3.36<sup>A</sup> | 30.4 ± 5.2<sup>A,B</sup> | 108.6 ± 9.2<sup>C</sup> | 87.2 ± 8.8<sup>B,C</sup> |
| **Fe<sub>a</sub>** [mg kg<sup>-1</sup>] | 1.66 ± 0.19<sup>A,B</sup> | 1.38 ± 0.58<sup>A</sup> | 10.7 ± 1.8<sup>B,C</sup> | 15.9 ± 1.1<sup>C</sup> |
| **K<sub>a</sub>** [mg kg<sup>-1</sup>] | 86.6 ± 16.1<sup>A</sup> | 142 ± 15<sup>A,B</sup> | 250 ± 22<sup>B</sup> | 251 ± 28<sup>B</sup> |
| **Ca<sub>a</sub>** [mg kg<sup>-1</sup>] | 1924 ± 60<sup>A</sup> | 2232 ± 91<sup>A,B</sup> | 2558 ± 113<sup>B</sup> | 2625 ± 25<sup>B</sup> |

### Evaluation of the CS severity

Potato tubers from each plant were washed with distilled water and CS severity was evaluated visually based on a fraction of tuber surface covered by lesions according to the standard area diagram proposed by Andrade et al. (43).

### Soil chemical analysis and respiration activity

Soil reaction (pH) was measured after 2 hours in a suspension of 2 g of soil in 5 mL of distilled H_2_O. Total content of C, N, and S in the soil was determined from 2 grams of homogenized soil samples from both the bulk at the start and the tuberosphere in the end of the experiment, which were dried, ground, and analyzed using a Vario MAX CNS analyzer (Elementar Analysensysteme, Hanau, Germany). The available fractions of soil potassium (K), calcium (Ca), iron (Fe) and phosphorus (P) contents were extracted with 1 M ammonium acetate (values are distinguished by ‘a’ in subscript) or 50 mM EDTA (‘e’ in subscript) (both 20 g soil in 100 mL) and assessed by optical emission spectroscopy with inductively coupled plasma (ICP-OES) by Aquatest (Prague, Czech Republic). Respiration activity of the soil community was measured in 5 g of sieved and homogenized soil by a pressure sensor method using OxiTop system with pressure sensor-data logger OxiTop-C heads including a capsule for NaOH solution (WTW, Weilheim, Germany) as described previously (44). The respiration rate was calculated from linear pressure decrease equivalent to O_2_ consumption between 50 and 250 h of incubation.

### DNA extraction, PCR amplification and sequencing

DNA was extracted from 0.5 g of soil samples according to Sagova-Mareckova et al. (45). The method is based on bead beating and phenol/chloroform extraction, followed by purification using CaCl_2_ and finally cleaned with GeneClean Turbo Kit (MP Biomedicals, Santa Ana, CA, USA). DNA extracts were stored at −70 °C. Fragments of the bacterial 16S rRNA gene including the variable region V4 were amplified by PCR using universal primers with 5’-linkers CS1_515F, 5’-ACA CTG ACG ACA TGG TTC TAC AGT GCC AGC MGC CGC GGT AA-3’, and CS2_806R 5’-TAC GGT AGC AGA GAC TTG GTC TGG ACT ACH VGG GTW TCT AAT-3’ (46). PCRs were performed in 25 μl reaction volumes using the Ex Taq HS DNA Polymerase (Takara, Kusatsu, Japan), and the PCR conditions were as follows: 5 min initial denaturation at 95 °C, followed by 28 cycles of: 30 s denaturation at 95 °C, 45 s annealing at 55 °C and 30 s extension at 72 °C. Construction of amplicon libraries including the second PCR and sequencing using Illumina MiSeq sequencer (Illumina, San Diego, CA) were done at the DNA Services Facility, Research Resources Center, University of Illinois (Chicago, IL). The Illumina MiSeq amplicon sequences of 16S rRNA genes are available in the NCBI Sequence Read Archive (www.ncbi.nlm.nih.gov/sra) as BioProject PRJNA1073595.

### Analysis of carbon, nitrogen, and phosphorus in potato leaves

Total carbon (C) and nitrogen (N) contents were determined in 2 g of dried potato leaves by high-temperature combustion followed by gas separation in a Vario MAX CNS analyzer by Aquatest. Total phosphorus (P) content was determined colorimetrically in 20 mg of dried potato leaves sequentially decomposed with HNO_3_ and HClO_4_ using the ammonium molybdate-ascorbic acid method on a QuikChem 8500 flow injection analytical system (Lachat Instruments, Hach Company, Loveland, CO, USA) in the Botanic institute in Třebon, Czech Republic (47).

### Data analysis

The raw sequence data were processed and analyzed in RStudio v. 2024.04.2 (48) with the R software environment v. 4.3.3 (49) utilizing DADA2 v. 1.28.0 (50) to inspect quality profiles, filter, and trim sequences and then infer amplicon sequence variants (ASVs) and remove chimeras. The resulting amplified sequence variants were classified in Mothur v. 1.47.0 software (51) using the SILVA Small Subunit rRNA Database, release 138.2 adapted for use in Mothur (https://mothur.s3.us-east-2.amazonaws.com/wiki/silva.nr_v138_2.tgz) as the reference database. ASVs of plastids and mitochondria, and those not classified in the domain *Bacteria* were removed from the ASV table. The names of higher taxa presented in the text were corrected to conform to the current taxonomy. The relative abundances of prokaryotic taxa were calculated using the *phyloseq* package (52) in RStudio and barplots were generated using the *ggplot2* package in RStudio. Non-metric multidimensional scaling (NMDS) was done using the *Mass* package based on Bray-Curtis distance matrices, and vectors of linear variables were fitted to the NMDS using function envfit from the package *vegan* (53) in RStudio. Metastats analysis (54) was performed in Mothur v. 1.47.0 software to show the numbers of ASVs that contributed significantly to the difference between prokaryotic communities in soils with different treatments. Differences in soil chemical and biological variables between treatments and sites, and nutrients contents in potato leaves were assessed by Kruskal-Wallis test with a post-hoc pairwise comparison using Dunn’s Test with Benjamini-Hochberg adjustment. Putative pools of functional genes in the communities were estimated based on the 16S rRNA gene sequence libraries using PICRUSt2 v.2.4.1 (55). This approach was used only to complement significant results by other more reliable methodology. Figures were edited using Inkscape (http://www.inkscape.org) and Microsoft Excel.

## Acknowledgements

The funding was obtained from the Ministry of Agriculture of the Czech Republic, Institutional Project MZE-RO0423 and project QL24010220. A.M conducted DNA extractions, PCRs, respective analyses and wrote a major part of MS, JK did experimental design, statistical analyses and created the more complicated figures, VT did all soil analyses, FM supplied the biochar including selections and analyses, MSM conceptualized the research, collected samples, acquired financial support, and finalized the MS.

